# EEG-triggered TMS reveals stronger brain state-dependent modulation of motor evoked potentials at weaker stimulation intensities

**DOI:** 10.1101/251363

**Authors:** Natalie Schaworonkow, Jochen Triesch, Ulf Ziemann, Christoph Zrenner

## Abstract

**Background:** Corticospinal excitability depends on the current brain state. The recent development of real-time EEG-triggered transcranial magnetic stimulation (EEG-TMS) allows studying this relationship in a causal fashion. Specifically, it has been shown that corticospinal excitability is higher during the scalp surface negative EEG peak compared to the positive peak of *µ*-oscillations in sensorimotor cortex, as indexed by larger motor evoked potentials (MEPs) for fixed stimulation intensity.

**Objective:** We further characterize the effect of *µ*-rhythm phase on the MEP input-output (IO) curve by measuring the degree of excitability modulation across a range of stimulation intensities. We furthermore seek to optimize stimulation parameters to enable discrimination of functionally relevant EEG-defined brain states.

**Methods:** A real-time EEG-TMS system was used to trigger MEPs during instantaneous brain-states corresponding to *µ*-rhythm surface positive and negative peaks with five different stimulation intensities covering an individually calibrated MEP IO curve in 15 healthy participants.

**Results:** MEP amplitude is modulated by *µ*-phase across a wide range of stimulation intensities, with larger MEPs at the surface negative peak. The largest relative MEP-modulation was observed for weak intensities, the largest absolute MEP-modulation for intermediate intensities. These results indicate a leftward shift of the MEP IO curve during the *µ*-rhythm negative peak.

**Conclusion:** The choice of stimulation intensity influences the observed degree of corticospinal excitability modulation by *µ*-phase. Lower stimulation intensities enable more efficient differentiation of EEG *µ*-phase-defined brain states.

## Introduction

The brain is ever active, with a rich dynamic structure of ongoing activity, even in the absence of task-related behavior. One salient feature of neurophysiological recordings are pronounced oscillatory rhythms, but the functional implications of these oscillations are not yet clear [1, 2]. A characterization of relevant brain states is challenging: what part of the signal is essential and what part is incidental? Additionally, how can we determine that a putative state-signature is functionally relevant? One promising approach uses brain-state triggered brain-stimulation to assess whether different EEG-derived state-signatures at the time of stimulation lead to different evoked potentials. Based on the hypothesis that oscillations organize cortical responses [3–5], the goal is to understand how different activity states lead to different functional consequences. In addition to providing a deeper understanding of the large-scale mesoscopic organization of the brain, this would allow selecting the optimal brain states for eliciting the desired behavioral responses through causal intervention, for instance by non-invasive brain stimulation.

Oscillatory rhythms have been shown to modulate cortical processing and influence perception and behavior. For instance, the oscillatory phase of the sensorimotor rhythms has been shown to change perceptual thresholds and behavioral responses [6–10] using correlative approaches. For transcranial magnetic stimulation (TMS), *post-hoc* estimation of EEG-defined brain-state at the time of stimulation is problematic, because the large stimulation artefact prevents use of standard signal processing methods (e.g. band-pass filtering) which require a window of data both before and after the time point of interest. Additionally, *post-hoc* evaluation of motor evoked potential (MEP) amplitude modulation by EEG phase requires a substantial number of trials per phase bin to achieve sufficient statistical power due to the well-known large inter-trial variability of MEP amplitudes [11, 12].

Real-time EEG-triggered TMS enables the functional consequences of different brain states to be probed in a causal manner and increases statistical power by preferentially targeting specific oscillatory phases. In the context of the motor system, a recent study [13] demonstrated a dependence of corticospinal excitability and plasticity on the phase of the cortical *µ*-rhythm using a real-time triggered EEG-TMS system. Larger MEP amplitudes were elicited by stimulation at time of *µ*-rhythm surface negative peak (N) compared to *µ*-rhythm positive peak (P). In that study, a fixed stimulation intensity (eliciting MEPs of on average of 1 mV peak-to-peak amplitude or using a fixed stimulus intensity of 120% of MEP threshold) was used to examine the effects of ongoing brain activity on corticospinal excitability.

The present study is motivated by the belief that the identification and characterization of functionally relevant EEG-defined large-scale brain-states is of critical importance for the development of more stable and effective personalized EEG-modulated therapeutic brain-stimulation protocols. The goal is to investigate the conditions under which functionally differentiable brain-states can be optimally identified in EEG-triggered TMS, specifically with regard to stimulus intensity.

Our recent computational modelling work suggests a larger relative excitability modulation by phase for lower stimulation intensities [14]. Here, we experimentally addressed the question of which stimulation parameters are optimal for the differentiation of *µ*-rhythm derived brain states. We investigated how *µ*-phase-modulation of corticospinal excitability changes as a function of stimulation intensity. Using a real-time EEGTMS set-up, pulses of five different stimulation intensities were triggered at two different oscillatory phase states (positive and negative peak) of the ongoing sensorimotor *µ*-rhythm, while MEPs were obtained to measure corticospinal excitability in each phase and intensity condition.

## Materials and Methods

### Experimental set-up

### EEG and EMG recordings

A combined EEG-TMS set-up was used to trigger stimulation pulses according to the instantaneous oscillatory phase of the recorded *µ*-rhythm. Scalp EEG was recorded from a 64-channel TMS compatible Ag/AgCl sintered ring electrode cap (EasyCap GmbH, Germany) in the international 10-20 system arrangement. Scalp electrode preparation consisted of light skin abrasion followed by filling with conductive gel (Electrode Cream, GE Medical Systems, USA) until an impedance of < 5 kΩ was reached. A 24-bit biosignal amplifier was used for combined 64-channel EEG and 2-channel EMG recordings (NeurOne Tesla with Digital Out Option, Bittium Biosignals Ltd., Finland), data was acquired in DC mode with a sample rate of 80 kHz at the head-stage and down-sampled online to a sample rate of 5 kHz. EMG was recorded from relaxed right abductor pollicis brevis (APB) and first dorsal interosseous (FDI) muscle with bipolar adhesive hydrogel electrodes (Kendall, Covidien) in a belly-tendon montage.

### TMS set-up

A passively cooled TMS double coil (PMD70-pCool, 70 mm winding diameter, MAG & More GmbH, Germany) was used together with a magnetic stimulator (Research 100, MAG & More GmbH, Germany) configured to deliver biphasic single cosine cycle pulses with 160 *µ*s period such that the second phase of the biphasic pulse induced an electrical field from lateral-posterior to medial-anterior, i.e., orthogonal to the central sulcus. Each TMS pulse was individually triggered through an external trigger input from the real-time system. Stimulation intensity was set programmatically using an analog control interface between 0–5 V and corresponding to 0–100% of maximum stimulator output through an analog output port interface (UEI PD2-MF-64-500/16L, United Electronic Instruments, USA) from the real-time system to allow for randomized ordering of intensity conditions. Stimulation was applied to the hand representation of left primary motor cortex (M1). The motor hot spot was identified as the coil position and orientation resulting consistently in maximum MEP amplitudes [15]. The target muscle was the muscle which responded to the lowest stimulator intensity and was then subsequently used to determine resting motor threshold (RMT) as the lowest intensity that elicited MEPs with a peak-to-peak amplitude of at least 50 *µ*V in 5 out of 10 trials [16]. A neuronavigation system (Localite GmbH, Sankt Augustin, Germany) was used to mark the coil position over the motor hot spot to monitor coil stability over time.

### Real-time EEG-triggered brain stimulation

The real-time processing system used in this experiment is described in detail in Zrenner et al. [13]. Briefly, an algorithm implemented in Simulink Real-Time (Mathworks Ltd, USA, R2016a) was used for real-time data acquisition, data processing and as the TMS stimulator control system. The algorithm was executed on an xPC Target processor (DFI-ACP CL630-CRM mainboard), processing online EEG data streamed through a real-time ethernet interface from the Digital Out interface of the EEG main unit in data packets at a rate of 1 kHz. The EEG signal used for real-time triggering was comprised of EEG channels overlying left sensorimotor cortex (C3, CP1, CP5, FC1, FC5), which were combined in a C3-centered Laplacian montage [17]. Data was down-sampled to 1 kHz by averaging and a sliding window of data of 500 ms width was used to compute estimates of instantaneous phase. The signal window was bandpass filtered (finite impulse response filter with order 128 and pass band 8–12 Hz), the last 64 ms were discarded because of edge artefacts and then forward predicted by an autoregressive model (Yule-Walker, order 30) for 128 ms. The analytic signal was computed by fast Fourier transform-based Hilbert transform, which was used to determine the instantaneous phase. In addition, the power spectrum was calculated from a sliding window of 1024 samples using Hann-windowed FFT and integrating spectral power in the 8–12 Hz frequency band. A digital output signal was generated to trigger the magnetic stimulator when three conditions were met: (1) an interstimulus interval (ISI) to the preceding pulse larger than 1.5 s (2) the temporal evolution of the phase estimate crosses the target phase. (3) a predefined *µ*-power threshold is met. The power-threshold was adjusted on an individual participant basis at the beginning of the experiment such that a median ISI of 2 seconds resulted. In the case of random-phase stimulation, only the minimum ISI and *µ*-power were considered as conditions for triggering the magnetic pulse, and a random delay was imposed between 0 and 100 ms.

### Experimental session

The experimental session consists of three types of blocks: (1) Resting state EEG recordings were made, with eyes open and closed (five minutes eyes open, participants instructed to fixate cross 1 meter in front of participant, followed by one minute eyes closed). (2) An IO curve was obtained (Fig. 1a). Eight intensities (90% to 160% RMT, in steps of 10%) were tested in randomized order, with 40 pulses per condition, applied with random-phase stimulation. The median ISI was 1.92 ± 0.24 s across participants. The IO curve was fitted with a logistic function *f*(*s*) = 1/(1 + exp(−*b* · (*s* − *a*))). Five stimulation intensities were chosen on the indvidual IO curve where median MEP amplitudes resulted at the following percentages of the maximum saturation level: SI_10% max_ / SI_30% max_/ SI_50% max_ / SI_70% max_ / SI_90% max_ (Fig. 1b). (3) *µ*-phase-triggered stimulation with real-time EEG-triggered TMS at surface *µ*-positive peak and surface *µ*-negative peak conditions was performed at the five intensities chosen in the previous block in randomized order with 60 pulses per condition (Fig. 1c). The phase-dependent stimulation block was repeated three times, i.e., 2 phases × 5 intensities × 60 pulses × 3 blocks = 1800 pulses total. The stimulator intensity was automatically adjusted between pulses, intensity conditions were applied in randomized order. The median ISI was 1.92 ± 0.15 s across participants. As an output measure, EMG was recorded (Fig. 1d).

**Figure 1:**
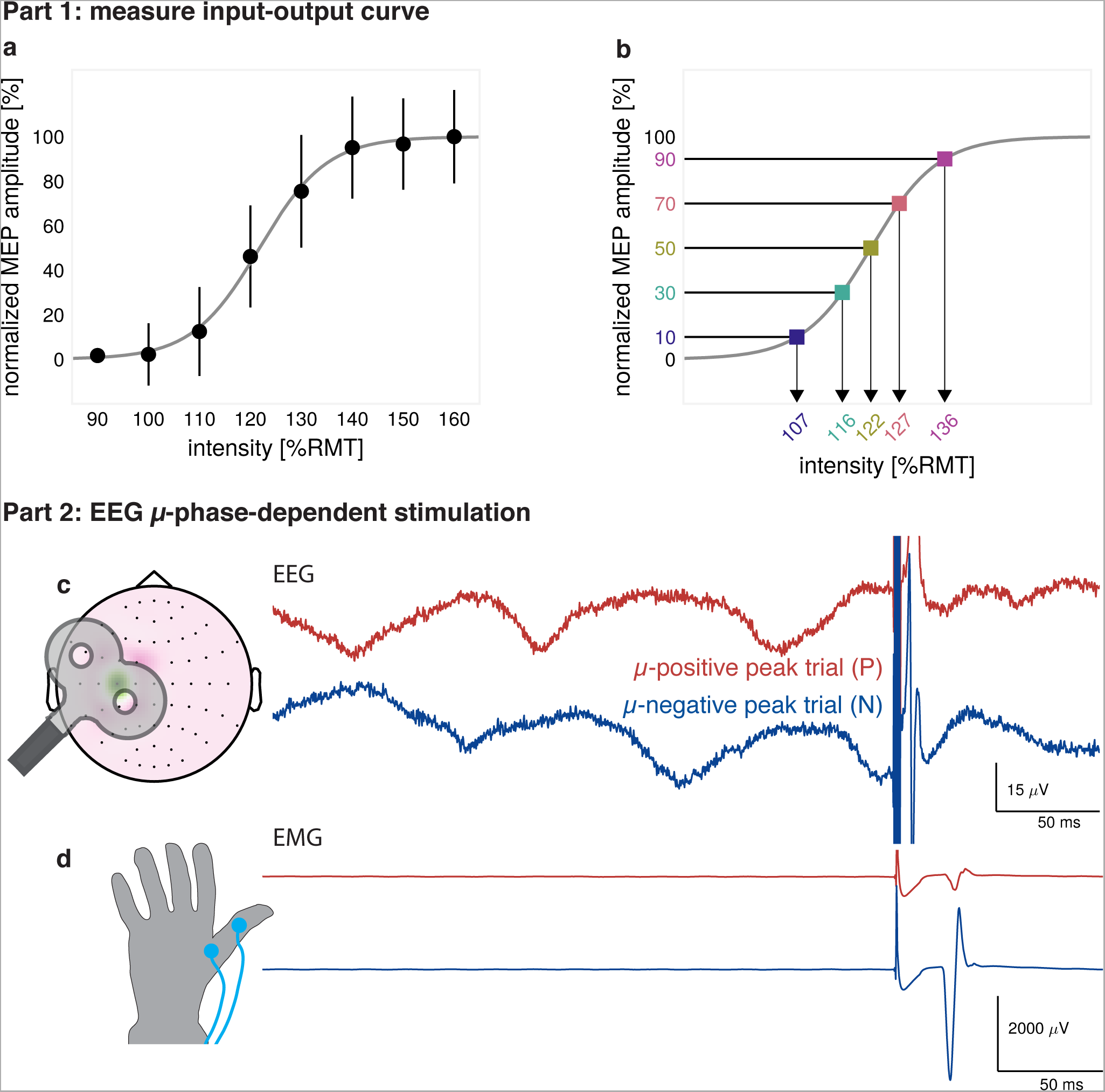
**Experimental Session**. (a) An IO curve is determined. Shown is an example IO curve for a single participant. A logistic function (grey curve) is fitted on median MEPs (black circles). Error bars indicate ± 1 SD. (b) From the logistic fit, five intensities (colored squares) are chosen for phase-dependent stimulation. (c) EEG *µ*-phase-triggered stimulation is performed, according to the instantaneous phase from the Laplace-filtered C3-signal. Two example single trials for the two types of trigger conditions, surface positive *µ*-peak (P) and surface negative *µ*-peak (N). (d) The corresponding single-trial EMG signals recorded for the two trigger conditions, with MEPs in the interval 20–40 ms after stimulation.

### Participants

The study protocol conformed to the Declaration of Helsinki and was approved by the local ethics committee at the medical faculty of the University of Tübingen (protocol 716/2014BO2). Written informed consent was obtained from all participants prior to the experiment. 17 right-handed participants (5 male, 12 female, mean age: 25.4 ± 2.6 years, age range: 22–32, average laterality score in Edinburgh handedness survey: 0.90 ± 0.12) with no history of neurological disease and usage of CNS drugs were selected according to the following inclusion criteria: (1) RMT of right FDI or APB muscle <= 62.5% of maximum stimulator output (MSO), so that a stimulation intensity range of up to 160% MSO (1.6*62.5%=100% MSO) could be explored. (2) The presence of a *µ*-rhythm with sufficient signal-to-noise ratio (SNR), as an adequate SNR is required for the phase-detection algorithm to estimate phases with sufficient accuracy. SNR was evaluated as follows (similar to Nikulin and Brismar [18]): A power function (*c* · *x^α^*) was fitted to the 1/f noise of the resting EEG data power spectrum from Laplace-filtered C3-electrode of each individual participant. For that, data points from frequency bins with typically no oscillatory components present (0.5–8 Hz, 30–40 Hz) were used. The fitted noise was subtracted from the power spectrum. The adjusted power in the 8–13 Hz band was assessed for a clearly identifiable peak in the *µ*-range, with 5 dB over noise level as inclusion threshold. Two participants were excluded from the experiment, one because of excessive pre-innervation in the EMG (with 54.1% of trials discarded according to the predefined threshold criterion), the other because the MEPs evoked by the fitted intensities for phase-dependent stimulation differed greatly (by 98.4%) from the IO curve fitted on median MEPs recorded in the pre-experiment, likely due to a coil position mismatch. This resulted in a sample size of 15 participants. Experiments were performed in accordance with current TMS safety guidelines [19]. All participants tolerated the procedures without any adverse effects.

### Data analysis and statistics

Data was analyzed with Matlab (Mathworks Ltd, USA, R2017b) and the BBCI toolbox [20]. EMG signals were high-pass filtered (Butterworth, filter order 2, 10 Hz). Trials with muscle activity 500 ms before stimulation onset were discarded with a threshold criterion (max-min amplitude > 50*µ*V). Peak-to-peak MEP amplitudes were determined within the interval of 20–60 ms after TMS pulse. All MEP time courses were inspected manually and non-response amplitudes were set to zero. Trials of the three phase-dependent blocks were pooled. For one participant, the last phase-dependent block was discarded because of excessive coil drift. The C3-centered EEG-Laplace filter extracted *µ*-rhythm pre-stimulation activity (500 ms before stimulation) was manually inspected for artefacts, and corresponding trials were discarded. Overall, 7.8% of trials were discarded for the phase-dependent stimulation sessions. We evaluated the relative as well as the absolute MEP differences between Nand P-trials. We quantified the relative phase-modulation by calculating the ratio 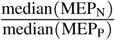 and the absolute phase-modulation by calculating 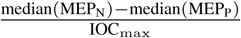 for each intensity condition, respectively, where IOC_max_ is the median MEP evoked at intensity 160% RMT, measuring the individual IO curve saturation level. We assessed the effect of intensity on phase-modulation by bootstrapping. Trials were randomly partitioned (with replacement) into two classes and the ratio and difference measures were calculated. This procedure was repeated for 100 000 iterations to arrive at confidence bounds and p-values.

## Results

### Methodological efficacy

To estimate the accuracy of the real-time phase-trigger algorithm, we determined the instantaneous phase by passing the five minutes resting EEG through the Simulink model from the experimental session to determine time points at which the algorithm would trigger. This procedure was chosen to avoid contamination by stimulation artefacts. Instantaneous phase was estimated by using Hilbert transform on the Laplacian C3 signal and band-pass filtered in 8–13 Hz frequency range. Phase prediction accuracy (mean ± standard deviation) across participants was −1.74° ± 53.65° in the positive peak condition and 178.26° ± 55.67° in the negative peak condition. Angular phase accuracy distribution plots for individual participants are shown in supplementary Fig. S1. The achieved phase accuracy was as expected similar to Zrenner et al. [13], as no changes were made to the core phase-detection algorithm.

To validate that the intensities chosen for the phase-triggered measurement session matched the section of the IO curve measured during the random-phase pre-measurement, the resulting averaged MEP amplitudes were compared by computing the percentile rank of the median MEP-amplitude for phase-stimulation conditions, pooled across N- and P-trials, assuming median MEPs from random-phase stimulation reflect an average of N- and P-trials. We found a mean deviation of −0.2% ± 12.0%, which allows adequate comparisons to the random-phase IO curve. The IO curves for individual participants can be found in supplementary Fig. S1e.

### Phase-modulation across stimulation intensity range

To illustrate the effect of *µ*-phase on MEP amplitudes, Fig. 2 shows data from one measurement session for an illustrative single participant, including the phase-dependent IO curves as well as MEP histograms, and the IO curve from the pre-experiment with random-phase stimulation. To quantify the effect of pre-stimulation *µ*-phase, we compared relative and absolute differences between MEP amplitudes of N- and P-trials.

**Figure 2:**
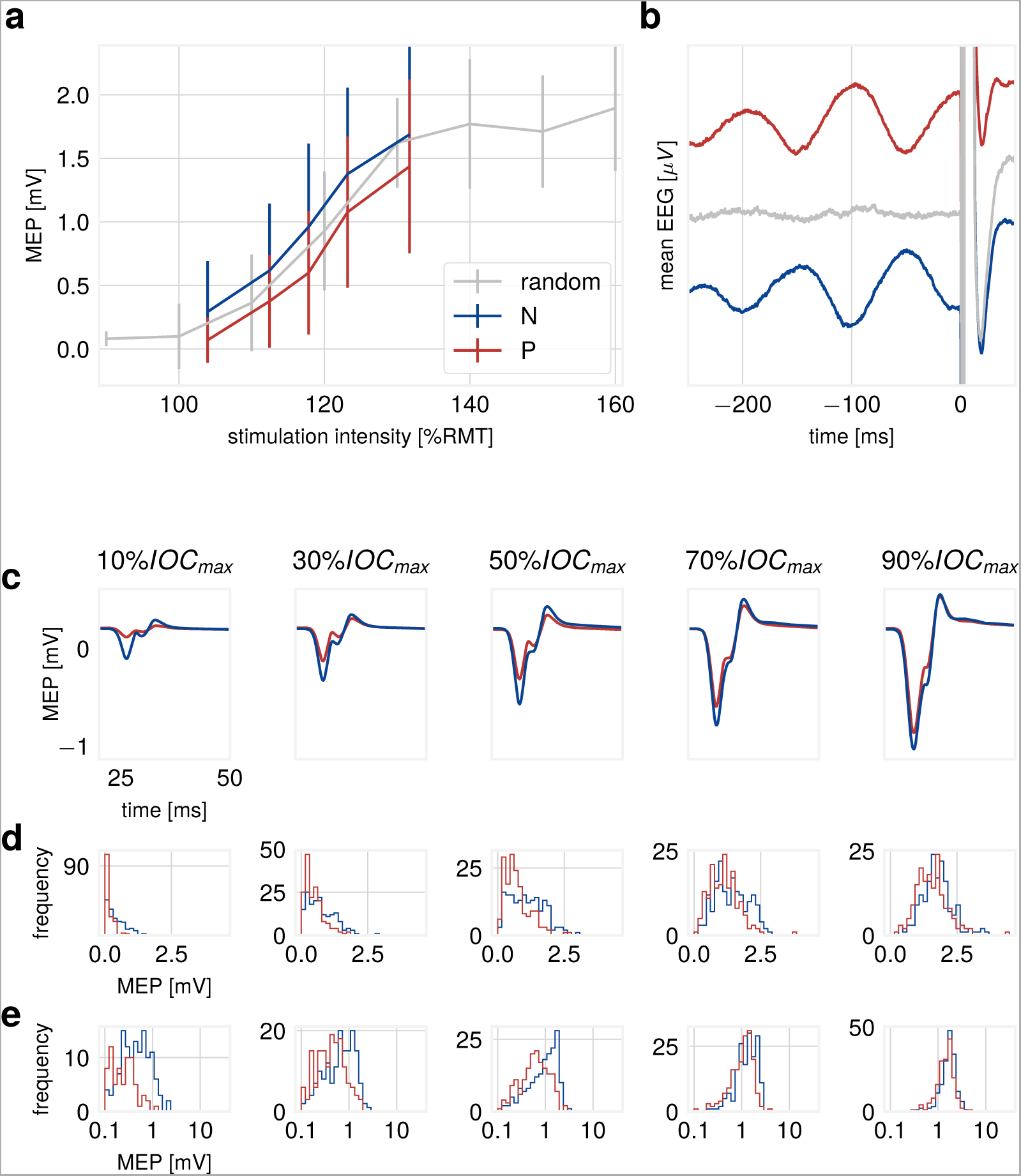
**Results for illustrative participant.** (a) IO curve for phase-dependent stimulation. Stimulation at the *µ*-phase negative peak (N) evokes MEPs of larger amplitudes (blue curve) than stimulation at the *µ*-phase positive peak (P, red curve). The random-phase IO curve runs in between (grey curve). Error bars indicate ± 1 SD. (b) Mean pre-stimulus C3 Laplacian-filtered EEG for N- and P-trials. (c) Mean MEP amplitudes (in mV) for the five stimulation intensities (blue: N-trials, red: P-trials) (d) Frequency distribution histograms of MEP amplitudes for the five stimulation intensity conditions on a linear scale (blue: N-trials, red: P-trials), non-responses excluded. (e) Histograms as in (d) but on a logarithmic scale to ease comparison at low intensity.

The mean influence of stimulation intensity on the degree of MEP amplitude modulation by *µ*-phase across all participants is shown in Fig. 3. We replicated the dependence of MEP amplitude on phase of the sensorimotor *µ*-rhythm at the time of TMS (i.e., larger MEP amplitudes at the negative peak of the *µ*-rhythm) [13] and detected a significant difference between N- and P-trials in four out of five intensity conditions. The effect of stimulation intensity on modulation by *µ*-phase was assessed as relative and absolute differences between N- and P-trials. The mean N/P-ratio decreased with higher intensity, (Fig. 3b). MEP amplitudes at the *µ*-negative peak were on average between 101% (at SI_10% max_, corresponding on average to 105% RMT, p = 0.006, two-tailed Wilcoxon signed-rank test) and 8% (at SI_90% max_, corresponding on average to 138% RMT, p>0.05) larger than MEPs evoked at the *µ*-positive peak. The N/P-difference peaked at the intermediate intensity SI_50% max_, with a group average difference between N- and P-trials of 10.5% of IOC_max_ (Fig. 3c). The measures for individual participants are shown in supplementary Figs. S2 and S3. For the participant with the largest observed N/P-ratio, the phase-dependent stimulation MEPs are lower compared to the random-phase stimulation IO curve (participant S16, supplementary Fig. S1e). This leads to increased N/P-ratio-values and to deviation from the N/P-ratio computed from the logistic fit (Fig. 3b). Variability of MEPs is higher at low intensities, as measured by the coefficient of variation (across participants, mean CV_10% max_=1.34, CV_30% max_=0.81, CV_50% max_=0.66, CV_70% max_=0.53, CV_90% max_=0.43). Due to higher variability of MEPs at low intensities, even large N/P-ratio values are not always significant within participants according to the bootstrap test.

**Figure 3:**
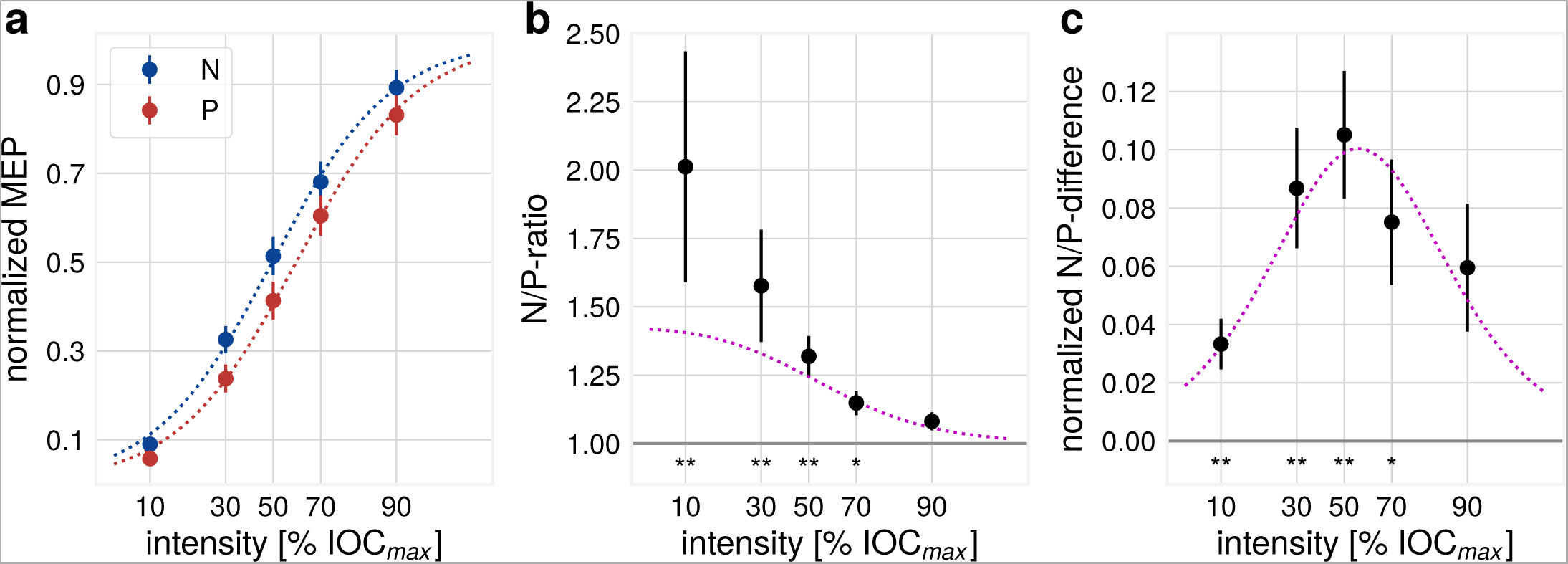
**Group level results.** (a) IO curve, for N- and P-trials, normalized to IOC_max_. A logistic function was fit to N- and P-trial median MEPs across intensity conditions separately (dotted lines). Error bars indicate ± 1 SEM. Dotted lines correspond to N/P-ratio values as calculated from the logistic fit. Error bars indicate ± 1 SEM. P-values (Bonferroni-corrected) for Wilcoxon signed-rank test across participants: **p ¡ 0.01 for SI_10% max_, SI_30% max_ and SI_50% max_, *p ¡ 0.05 for SI_70% max_ and p>0.05 for SI_90% max_. (c) N/P-difference between N- and P-trials. Normalized to IOC_max_, Dotted lines correspond to N/P-difference values as calculated from the logistic fit. Error bars indicate ± 1 SEM.

To illustrate the effect of the sample size on the probability of detecting a difference between N- and P-trials for every intensity condition we used a simulation approach. For each participant, MEPs were resampled with replacement separately for Nand P-trials for varying sample sizes and a Wilcoxon rank-sum test was performed, noting whether the null hypothesis was rejected. This procedure was repeated 10 000 times. The results are shown in Fig. 4.

**Figure 4:**
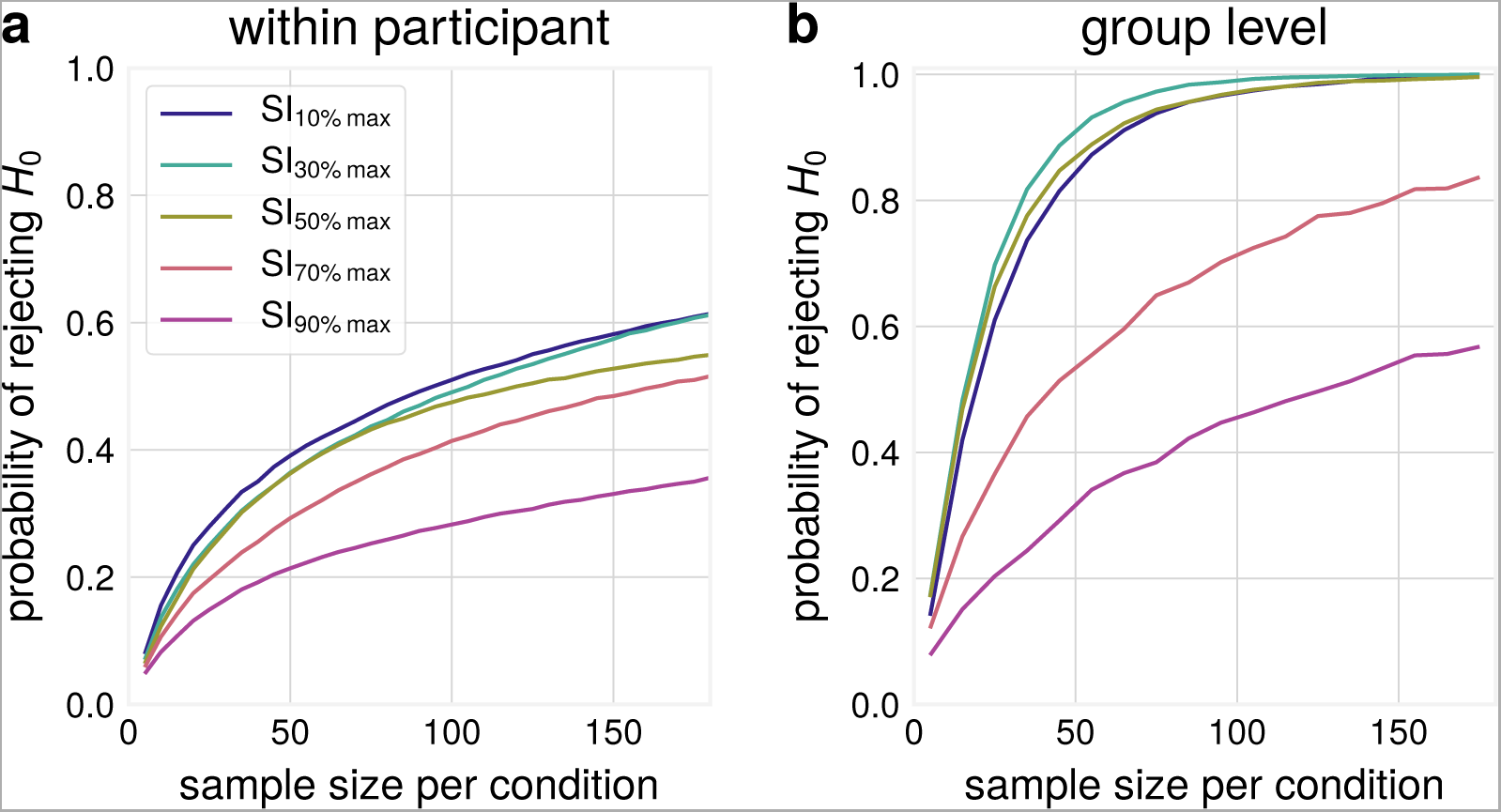
**Illustration of the effect of sample size on the probability of detecting a difference between N- and P-trials.** The dependence of the probability to reject the null hypothesis of no difference between N- and P-trials on the used sample size per condition for different stimulation intensities in (a) within participant comparisons and (b) group level comparisons (N=15 participants included in this study).

Using 180 trials per condition, the mean probability to reject the null hypothesis (N/P-ratio equals 1 or N/P-difference equals 0) within a single participant is 62% for the CV_10% max_ and CV_30% max_ conditions, and around 35% for the CV_90% max_ condition. Using only 90 trials, these values decline to 50% and 28%, respectively. This demonstrates that the required number of trials to differentiate between *µ*-phase-defined states with high statistical power within participants can be reduced by choosing a lower stimulation intensity. In this analysis, also non-responders (i.e. participants without a significant phase-modulation for any tested stimulation intensity) are included, as the goal of this analysis is to show relative differences in required sample sizes between intensities. For non-responders, the null hypothesis may be true, therefore the actual statistical power at a given sample size is higher than the average probability of rejecting *H*_0_. At the group level, using a low to intermediate intensity will enable detection of significant effects of the size observed in the current study with high certainty.

### Non-Responders

In our study, 7 of the 15 included participants did not show a significant MEP amplitude modulation by instantaneous *µ*-phase at any of the tested stimulation intensities, which is consistent with the data reported in Zrenner et al. [13]. We performed a number of correlation analyses in order to identify possible factors, which may separate responders from non-responders: We found no significant correlation between SNR as obtained from resting state EEG data and the effect size for SI_10% max_ (p>0.05, Spearman rank correlation). Overall, higher SNR resulted in improved performance of the phase-detection algorithm as measured by decreased standard deviation of the phase accuracy (r=-0.67, p=0.0081, Spearman rank correlation), but pronounced rhythm with high SNR did not translate to a larger phase-modulation effect. A significant sub-population of participants remained that showed a clear *µ*-rhythm without exhibiting a clear phase-modulatory effect on MEP amplitude. Additionally, no significant correlation was observed between distance of coil to center electrode of the Laplacian filter and effect size at SI_10% max_ (p>0.05, Spearman rank correlation). In this study, a standard MNI brain model was used for navigation. In future studies with individual participant MRI anatomy additional factors could be investigated, such as the orientation of dipoles underlying the mean topography around the stimulation trigger (supplementary Fig. S1b), coil position and orientation relative to the central sulcus. Therefore, the factors required for observing *µ*-phase modulation of corticospinal excitability remain to be further elucidated.

## Discussion

We replicated the finding that corticospinal excitability as measured by MEP amplitude is modulated by the phase of the ongoing *µ*-rhythm [13], with larger MEP amplitudes at the negative compared to the positive peak. Additionally, in agreement with predictions based on our modeling work [14], we demonstrated that the magnitude of the modulatory effect depends on stimulation intensity, with largest relative modulation for low intensities and largest absolute differences for intermediate stimulation intensities. The reduced modulatory influence of *µ*-rhythm at high stimulation intensities can be explained by saturation of the IO curves of MEPs. If stimulation intensity is sufficiently far above the motor threshold, any ongoing fluctuations of that threshold influence behavioral outcomes to a lesser degree and will result in smaller relative differences between N- and P-trials. This is compatible with many previous findings showing greatest sensitivity of MEP amplitude to intervention in the low and/or intermediate parts of the IO curve (e.g. [21–24]).

### Implications

One practical implication of the present findings is that studies seeking to demonstrate differential EEG-defined brain states based on differential TMS-evoked responses should be designed with a sufficient number of interleaved trials (more than 100 per condition) and use low stimulation intensity to maximize statistical power, where the lower limit of intensity is determined by the proportion of non-responses and measurement noise when quantifying small responses. Based on this study, the intensity setting resulting in an average MEP amplitude 20% of IOC_max_ would be recommended; in our dataset, this corresponded to a stimulus intensity of on average 112% RMT, with 10 of the 15 participants in our sample in the range between 108–117 %RMT. However, depending on the IO curve, a stimulation intensity based on a fixed RMT percentage can already be in the saturation range of an individual participant, where MEP amplitudes do not significantly differ between *µ*-phase conditions. Measuring the individual IO curve and specifically estimating the maximum amplitude at saturation is therefore helpful to choose an optimal intensity.

### Limitations

The range of TMS intensities used is limited at the lower end by the motor threshold, as stimulation at intensities significantly below RMT does not result reliably in EMG responses in any brain-state. Nevertheless, sub-threshold TMS does affect cortical circuits, as can be demonstrated, for instance, in paired-pulse paradigms (e.g., [25, 26]), with pre-innervation [27], or in TMS evoked EEG potentials [28, 29]. It is therefore feasible (using paired-pulse protocols, performing stimulation during pre-innervation, or using cortical EEG responses) to investigate in subsequent studies whether EEG-defined brain-states can be differentiated at intensities below single-pulse RMT.

In addition to phase, pre-stimulus oscillatory power has been shown to modulate perception [30–33], but in our experimental design, the impact of instantaneous *µ*-power (or power in other frequency bands) on MEP amplitude is difficult to assess. As we used a power-threshold, to ensure suitable accuracy of the phase-detection algorithm, no trials with low *µ*-power were acquired. Real-time triggered EEG-TMS could be used in the future to explore the role of instantaneous power.

Not all participants displayed a modulation of MEPs by instantaneous *µ*-phase. We tried to identify factors which predict the individual degree of phase-modulation. High SNR alone is not sufficient for a large effect, as in our data set, there are participants with high SNR but no modulation of MEP amplitude by *µ*-phase. Furthermore, the standard Laplacian C3 filter may not extract the functionally relevant oscillatory component, depending on individual anatomical features. Future studies may improve this aspect by using individualized spatial filters or anatomically guided source level analysis. This may also increase the proportion of participants which can be studied with phase-dependent brain stimulation, as in this study only participants with a *µ*-rhythm as detected by a standard C3 Hjorth montage were included, whereas individualized approaches will yield increased SNR for oscillatory components.

The conditions required to establish a clear dependence of cortical excitability on instantaneous *µ*-phase are not sufficiently understood yet. Understanding this dependence will yield a clearer view on what exactly is stimulated by TMS, and on the interplay of endogeneous oscillatory rhythms in motor areas and their functional role.

## Acknowledgements

We thank Anna Kempf and Tamara Vasilkovska for help with participant coordination and experimental preparation. NS and JT acknowledge support from the Quandt foundation. NS and CZ are supported through a German Federal Ministry for Economic Affairs and Energy of EXIST Transfer of Research Grant. CZ acknowledges support from the Clinician Scientist Program at the Faculty of Medicine at the University of Tübingen. UZ acknowledges support from the German Research Foundation (DFG, grant ZI 542/7-1). This research was supported with funding from the Federal Ministry of Education and Research of Germany through the MOTOR-BIC project (BMBF, grant 13GW0053A).

## Supplementary Material

**Figure S1:**
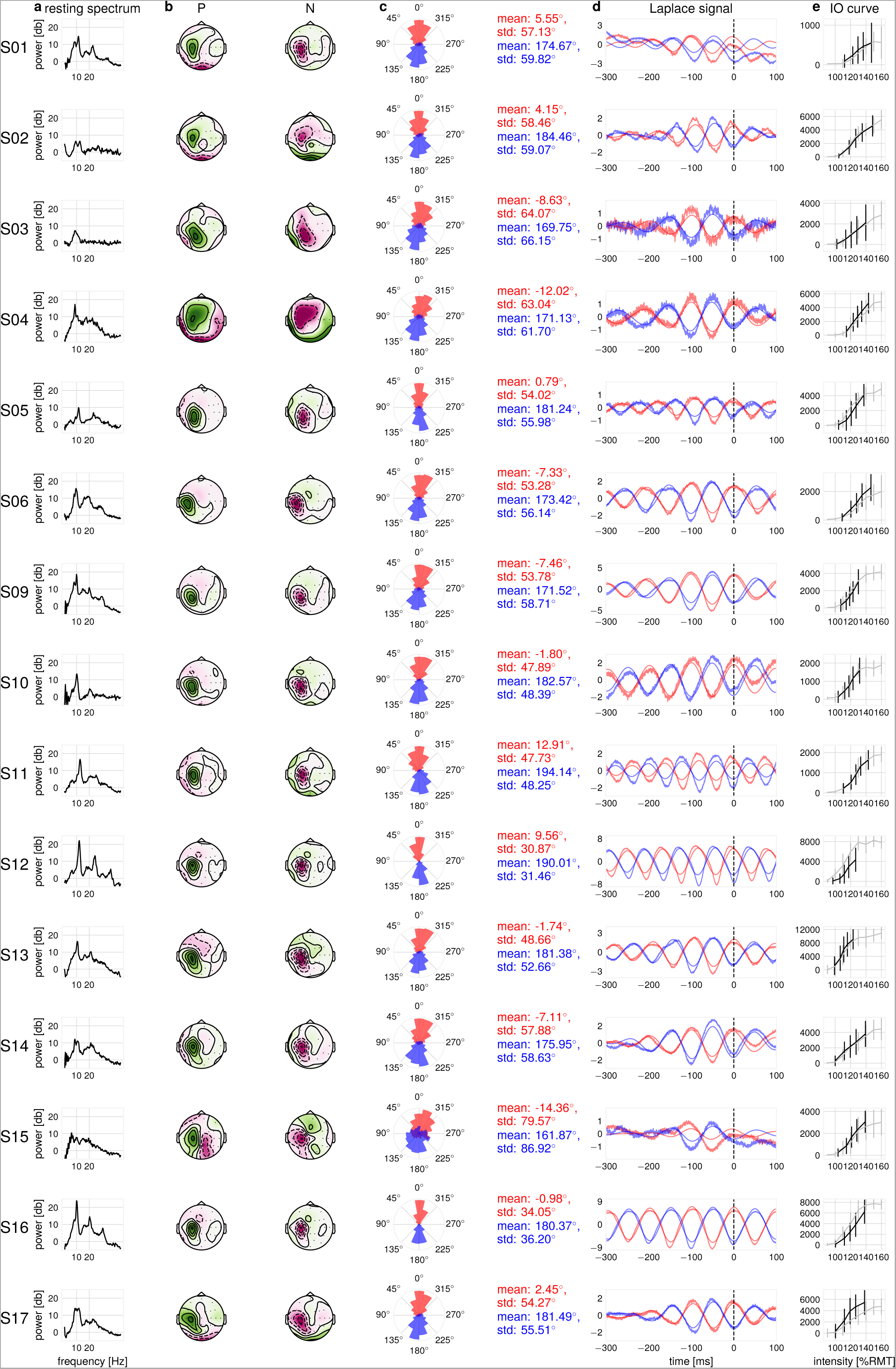
**Methodological accuracy data of all included individual participants.** Participants S07 and S08 were excluded from analysis. Each row shows the data for one participant. (a) The adjusted resting state EEG spectrum for evaluating SNR, Hann window of 16 s. (b) Mean Laplacian topography [-10 10] ms around the non-stimulated positive (P) and negative (N) triggers, color scale individually adjusted, with green signaling positive values and pink negative values. (c) Phase accuracy rose plots for N- and P-trials as computed for non-stimulated trials from resting state EEG data (d) Mean Laplacian C3 pre-stimulation activity for non-stimulated trials, raw and band-pass filtered (8–12 Hz). (e) IO curve, gray line indicates random phase IO curve from pre-experiment, in black, the phase-dependent stimulation trials, pooled across N- and P-trials.

**Figure S2:**
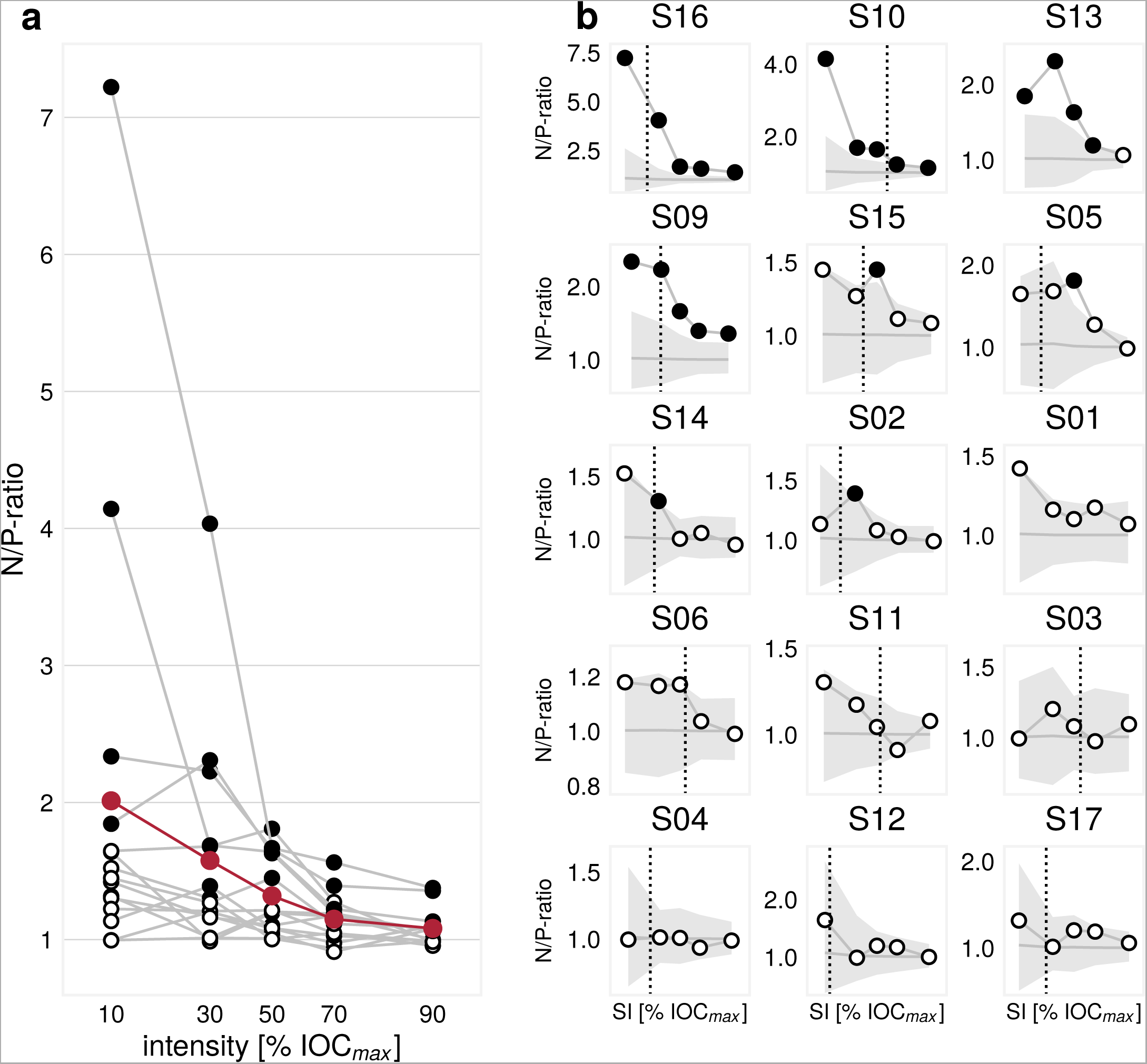
**N/P-ratio for individual participants** (a) N/P-ratio for individual participants, and mean (red curve). Filled circles represent N/P-ratio-values larger than the 99.5^th^ percentile of bootstrapped values. (b) N/P-ratio for all individual participants. The shaded area marks the 0.5^th^–99.5^th^ percentile range of bootstrapped N/P-ratio-values. The black dotted line indicates individual intensity producing 1 mV MEPs (if SI_1*mV*_ is within the range of the tested stimulation intensities).

**Figure S3:**
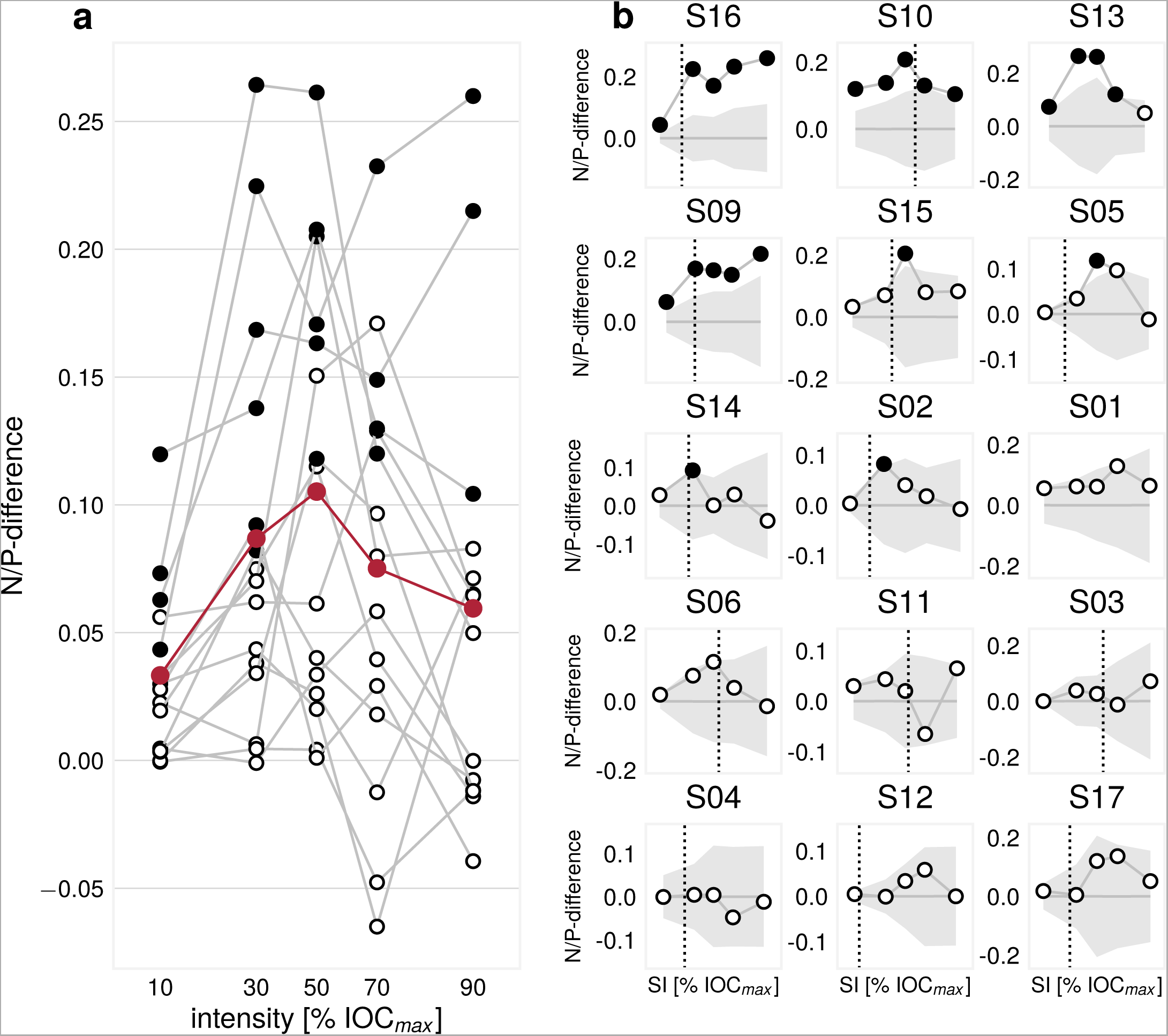
**N/P-difference for individual participants**. (a) N/P-difference for individual participants, and mean (red curve). Filled circles represent difference-values larger than the 99.5^th^ percentile of bootstrapped N/P-difference-values. (b) N/P-difference for all individual participants. The shaded area marks the 0.5^th^–99.5^th^ percentile range of bootstrapped N/P-difference-values. The black dotted line indicates individual intensity producing 1 mV MEPs (if SI_1*mV*_ is within the range of the tested stimulation intensities).

## References

[1] Wang XJ. Neurophysiological and Computational Principles of Cortical Rhythms in Cognition. Physiol Rev 2010;90(3):1195–268.

[2] Buzsáki G, Draguhn A. Neuronal oscillations in cortical networks. Science 2004;304(5679):1926–9.

[3] VanRullen R, Busch NA, Drewes J, Dubois J. Ongoing EEG phase as a trial-by-trial predictor of perceptual and attentional variability. Front Psychol 2011;2(60):1–9.

[4] Fries P. A mechanism for cognitive dynamics: Neuronal communication through neuronal coherence. Trends Cogn Sci 2005;9(10):474–80.

[5] Jensen O, Mazaheri A. Shaping Functional Architecture by Oscillatory Alpha Activity: Gating by Inhibition. Front Hum Neurosci 2010;4(4):1–8.

[6] Busch NA, Dubois J, VanRullen R. The phase of ongoing EEG oscillations predicts visual perception. J Neurosci 2009;29(24):7869–76.

[7] Mathewson KE, Gratton G, Fabiani M, Beck DM, Ro T. To see or not to see: prestimulus phase predicts visual awareness. J Neurosci 2009;29(9):2725–32.

[8] Dugué L, Marque P, VanRullen R. The phase of ongoing oscillations mediates the causal relation between brain excitation and visual perception. J Neurosci 2011;31(33):11889–93.

[9] Ai L, Ro T. The phase of prestimulus alpha oscillations affects tactile perception. J Neurophysiol 2013;111(6):1300–7.

[10] Keil J, Timm J, SanMiguel I, Schulz H, Obleser J, Schönwiesner M. Cortical brain states and corticospinal synchronization influence TMS-evoked motor potentials. J Neurophysiol 2014;111(3):513–9.

[11] Kiers L, Cros D, Chiappa K, Fang J. Variability of motor potentials evoked by transcranial magnetic stimulation. Electroen Clin Neuro 1993;89(6):415–23.

[12] Ellaway P, Davey N, Maskill D, Rawlinson S, Lewis H, Anissimova N. Variability in the amplitude of skeletal muscle responses to magnetic stimulation of the motor cortex in man. Electroen Clin Neuro 1998;109(2):104–13.

[13] Zrenner C, Desideri D, Belardinelli P, Ziemann U. Real-time EEG-defined excitability states determine efficacy of TMS-induced plasticity in human motor cortex. Brain Stimul 2017;(in press). doi:10.1016/j.brs.2017.11.016.

[14] Schaworonkow N, Triesch J. Ongoing brain rhythms shape I-wave properties in a computational model. bioRxiv 2017;:205450.

[15] Rossini P, Burke D, Chen R, Cohen LG, Daskalakis Z, Di Iorio R, et al. Non-invasive electrical and magnetic stimulation of the brain, spinal cord, roots and peripheral nerves: Basic principles and procedures for routine clinical and research application. An updated report from an I.F.C.N. Committee. Clin Neurophysiol 2015;126(6):1071–107.

[16] Groppa S, Oliviero A, Eisen A, Quartarone A, Cohen L, Mall V, et al. A practical guide to diagnostic transcranial magnetic stimulation: report of an IFCN committee. Clin Neurophysiol 2012;123(5):858–82.

[17] Hjorth B. An on-line transformation of EEG scalp potentials into orthogonal source derivations. Electroen Clin Neuro 1975;39(5):526–30.

[18] Nikulin VV, Brismar T. Phase synchronization between alpha and beta oscillations in the human electroencephalogram. Neurosci 2006;137(2):647–57.

[19] Rossi S, Hallett M, Rossini PM, Pascual-Leone A, of TMS Consensus Group S, et al. Safety, ethical considerations, and application guidelines for the use of transcranial magnetic stimulation in clinical practice and research. Clin neurophysiol 2009;120(12):2008–39.

[20] Blankertz B, Acqualagna L, D¨ahne S, Haufe S, Schultze-Kraft M, Sturm I, et al. The Berlin brain-computer interface: progress beyond communication and control. Front Neurosci 2016;10.

[21] Ikoma K, Samii A, Mercuri B, Wassermann E, Hallett M. Abnormal cortical motor excitability in dystonia. Neurology 1996;46(5):1371–.

[22] Devanne H, Lavoie B, Capaday C. Input-output properties and gain changes in the human corticospinal pathway. Exp Brain Res 1997;114(2):329–38.

[23] Di Lazzaro V, Oliviero A, Profice P, Pennisi M, Pilato F, Zito G, et al. Ketamine increases human motor cortex excitability to transcranial magnetic stimulation. J Physiol 2003;547(2):485–96.

[24] Lücke C, Heidegger T, Röhner M, Toennes SW, Krivanekova L, Müller-Dahlhaus F, et al. Deleterious effects of a low amount of ethanol on LTP-like plasticity in human cortex. Neuropsychopharmacol 2014;39(6):1508–18.

[25] Kujirai T, Caramia MD, Rothwell JC, Day BL, Thompson PD, Ferbert A, et al. Corticocortical inhibition in human motor cortex. J Physiol 1993;471:501–19.

[26] Ziemann U, Rothwell JC, Ridding MC. Interaction between intra-cortical inhibition and facilitation in human motor cortex. J Physiol 1996;496:873–81.

[27] Di Lazzaro V, Restuccia D, Oliviero A, Profice P, Ferrara A, Insola A, et al. Effects of voluntary contraction on descending volleys evoked by transcranial stimulation in conscious humans. J Physiol 1998;508(2):625–33.

[28] Komssi S, K¨ahkönen S, Ilmoniemi RJ. The Effect of Stimulus Intensity on Brain Responses Evoked by Transcranial Magnetic Stimulation. Hum Brain Mapp 2004;21(3):154–64.

[29] Fecchio M, Pigorini A, Comanducci A, Sarasso S, Casarotto S, Premoli I, et al. The spectral features of EEG responses to transcranial magnetic stimulation of the primary motor cortex depend on the amplitude of the motor evoked potentials. PloS one 2017;12(9):e0184910.

[30] Hanslmayr S, Aslan A, Staudigl T, Klimesch W, Herrmann CS, B¨auml KH. Prestimulus oscillations predict visual perception performance between and within subjects. Neuroimage 2007;37(4):1465–73.

[31] Van Dijk H, Schoffelen JM, Oostenveld R, Jensen O. Prestimulus oscillatory activity in the alpha band predicts visual discrimination ability. J Neurosci 2008;28(8):1816–23.

[32] Linkenkaer-Hansen K, Nikulin VV, Palva S, Ilmoniemi RJ, Palva JM. Prestimulus oscillations enhance psychophysical performance in humans. J Neurosci 2004;24(45):10186–90.

[33] Weisz N, Wühle A, Monittola G, Demarchi G, Frey J, Popov T, et al. Prestimulus oscillatory power and connectivity patterns predispose conscious somatosensory perception. Proc Natl Acad Sci 2014;111(4):E417–25.

